# Rapid evolution of genes with anti-cancer functions during the origins of large bodies and cancer resistance in elephants

**DOI:** 10.1101/2024.02.27.582135

**Authors:** Jacob Bowman, Vincent J. Lynch

## Abstract

Elephants have emerged as a model system to study the evolution of body size and cancer resistance because, despite their immense size, they have a very low prevalence of cancer. Previous studies have found that duplication of tumor suppressors at least partly contributes to the evolution of anti-cancer cellular phenotypes in elephants. Still, many other mechanisms must have contributed to their augmented cancer resistance. Here, we use a suite of codon-based maximum-likelihood methods and a dataset of 13,310 protein-coding gene alignments from 261 *Eutherian* mammals to identify positively selected and rapidly evolving elephant genes. We found 496 genes (3.73% of alignments tested) with statistically significant evidence for positive selection and 660 genes (4.96% of alignments tested) that likely evolved rapidly in elephants. Positively selected and rapidly evolving genes are statistically enriched in gene ontology terms and biological pathways related to regulated cell death mechanisms, DNA damage repair, cell cycle regulation, epidermal growth factor receptor (EGFR) signaling, and immune functions, particularly neutrophil granules and degranulation. All of these biological factors are plausibly related to the evolution of cancer resistance. Thus, these positively selected and rapidly evolving genes are promising candidates for genes contributing to elephant-specific traits, including the evolution of molecular and cellular characteristics that enhance cancer resistance.

## Introduction

Large body sizes have repeatedly evolved in many mammalian lineages such as whales (*Cetaceans*), elephants and their extinct relatives (*Proboscideans*), extinct hornless rhinos (*Paraceratherium*, aka ‘Walter’), giant ground sloths (*Megatherium*), and giant armadillos (*Glyptodon*); this tendency for lineages to increase in size over evolutionary time is relatively common across the tree of life (Purvis and Orme, 2005). There are many constraints on the evolution of gigantic body sizes, including an increased risk of developing cancer. For example, if all cells in all organisms have a similar risk of malignant transformation and equivalent cancer suppression mechanisms, then organisms with many cells should have a higher prevalence of cancer than organisms with fewer cells, particularly because the size of most cell types is similar across species with different body sizes (Hoogduijn et al., 2013; Savage et al., 2007). While there is a positive correlation between body size and cancer prevalence within species, for example, humans and dogs (Benyi et al., 2019; Dobson, 2013; Green et al., 2011; Zhou et al., 2022), there is no positive correlation between body size and cancer risk between mammalian species (Abegglen et al., 2015; Bulls et al., 2022; Compton et al., 2023); this lack of correlation is called ‘Peto’s Paradox’ (Peto, 2015, 1975). Indeed, within mammals, there is a statistically significant negative correlation between body mass and cancer prevalence (Bulls et al., 2022), indicating that mammals with enormous bodies have evolved particularly effective anticancer mechanisms.

*Afrotherian* mammals are an excellent model system to study the evolution of cancer resistance because while most lineages such as golden moles (∼70g), tenrecs (120g), elephant shrews (∼170g), hyraxes (∼3kg), aardvarks (∼60kg), and sea cows (600-900kg) are small- to large-bodied, gigantic *Proboscideans* including African Savannah (4,800kg) and Asian elephants (3,200kg) and extinct species such as *Deinotherium* (∼12,000kg), *Mammut borsoni* (16,000kg), and the straight-tusked elephant (∼14,000kg) are nested deeply within the small-bodied species and evolved from a small-bodied ancestor (Elliot and Mooers, 2014; Puttick and Thomas, 2015; Vazquez and Lynch, 2021). Large body masses in *Proboscideans* evolved stepwise (O’Leary et al., 2013a, 2013b; Puttick and Thomas, 2015; Springer et al., 2013; Vazquez and Lynch, 2021), coincident with dramatic reductions in intrinsic cancer risk (Vazquez and Lynch, 2021) and the evolution of anticancer cellular phenotypes. For example, elephant cells induce apoptosis at very low levels of DNA damage (Abegglen et al., 2015; Sulak et al., 2016a; Vazquez et al., 2018), are resistant to oxidative stress-induced cell death (Gomes et al., 2011), are resistant to experimental immortalization (Fukuda et al., 2016; Gomes et al., 2011), and repair DNA damage more rapidly than cells from smaller bodied species (Francis et al., 1981; Hart and Setlow, 1974; Promislow, 1994).

Many mechanisms must have contributed to the evolution of enhanced cancer resistance. For example, the sensitivity of elephant cells to DNA damage is mediated in part by an increase in the number of tumor suppressor genes in the elephant lineage (Caulin et al., 2015; Doherty and Magalhães, 2016; Sulak et al., 2016b; Tollis et al., 2020; Vazquez et al., 2018; Vazquez and Lynch, 2021) whereas the resistance to oxidative stress-induced cell death may be mediated by a duplicate *superoxide dismutase 1* (*SOD1*) gene (Vazquez and Lynch, 2021). Here, we explore the possibility that the functional divergence of protein-coding genes may also have contributed to anti-cancer traits in the elephant lineage by identifying genes with evidence of positive selection and rapid evolution, an evolutionary signature often associated with the functional divergence of proteins (Slodkowicz and Goldman, 2020; Tennessen, 2008). We used maximum likelihood models to characterize the strength and direction of selection acting on 13,310 orthologous coding gene alignments from the genomes of 261 *Eutherian* mammals and identified several hundred positively selected and rapidly evolving genes in the elephant lineage. These genes are enriched in gene ontology (GO) and pathway terms related to various kinds of regulated cell death, DNA damage repair, regulation of the cell cycle, EGFR–signaling, and immune functions, particularly neutrophil biology, which have essential roles in cancer biology (Hedrick and Malanchi, 2021; Huang et al., 2020; Mollinedo, 2019; Normanno et al., 2006; Sigismund et al., 2018; Vorobjeva and Chernyak, 2020). Thus, these are good candidates for genes that contribute to elephant-specific traits, including the evolution of cancer resistance.

## Results

### Hundreds of genes experienced an episode of positive selection or rapid evolution in elephants

Episodic adaptive evolution (“positive selection”) is thought to be associated with functional divergence (Slodkowicz and Goldman, 2020; Tennessen, 2008). Therefore, we used two complementary methods, ABSREL (Smith et al., 2015) and BUSTED (Murrell et al., 2015a), to identify genes with evidence of positive selection in the elephant lineage. We used a dataset of alignments from 13,310 orthologous genes from 261 *Eutherian* mammals that we previously reported (Bowman et al., 2023b) and a phylogeny with *Paenungulates* collapsed to a polytomy to reflect that most genes lack significant phylogenetic support for the resolution of this clade (**Figure 1A**) (Bowman et al., 2023a). The ABSREL and BUSTED models are variants of the branch-sites random effect likelihood (BSREL) model of coding sequence evolution (Pond et al., 2011a; Smith et al., 2015), which allows for variation in *d_N_*/*d_S_* (ω) rates across lineages and sites; however, while ABSREL selects the optimal number of rate categories for each gene, BUSTED imposes three rate categories such that ω_1_≤ω_2_≤ω_3_ (**Figure 1B**). Both models also accommodate synonymous rate variation across sites [S] (Pond and Muse, 2005; Wisotsky et al., 2020), double and triple multi-nucleotide mutations per codon [MH] (Lucaci et al., 2021), and both synonymous rate variation and multi-nucleotide mutations [SMH]; failure to account for synonymous rate variation (Dunn et al., 2019; Rahman et al., 2021; Wisotsky et al., 2020) and multi-nucleotide mutations can lead to widespread false positive inferences of positive selection (Dunn et al., 2019; Lucaci et al., 2023a; Nozawa et al., 2009; Suzuki, 2008; Venkat et al., 2018; Yang and Reis, 2011). We used small sample corrected Akaike Information Criterion values (ΔAICc≥10) to select for the best [S], [MH], and [SMH] model and inferred positive selection for a gene when the best fitting model included a class of sites with ω>1 and a likelihood ratio test *P*≤0.05.

**Figure 1.**
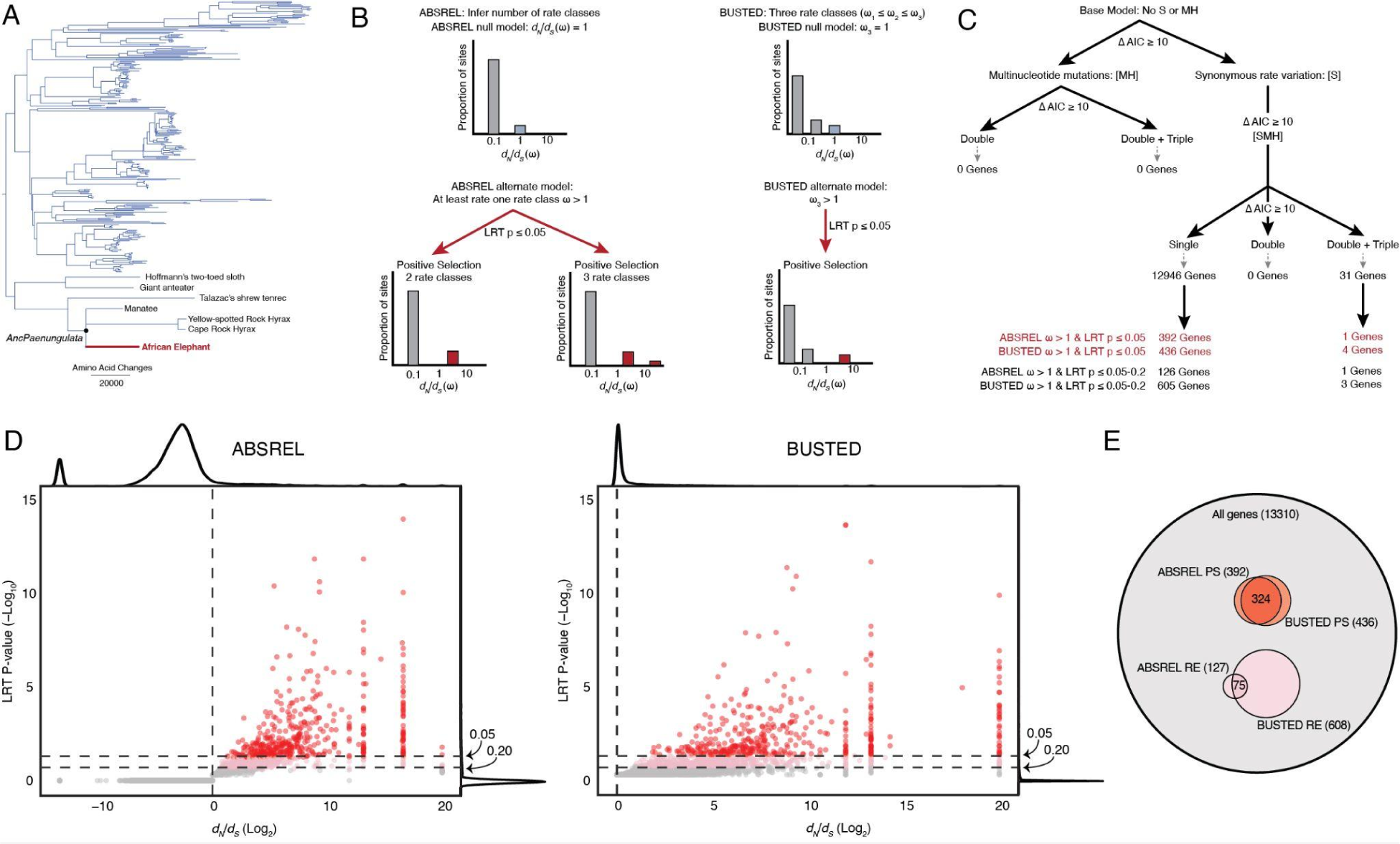
Identification and positively selected and rapidly evolving genes in the elephant lineage. **A.** Phylogeny of 261 Eutherian mammals used in selection tests. The Atlantogenatan part of the tree is expanded to show which species of Afrotheria and Xenarthra are included in the analyses. Branch lengths are shown as amino acid changes per lineage. **B.** Cartoon description of the ABSREL and BUSTED models. **C.** Decision tree showing [S], [MH], and [SMH] model selection (inferred with ABSREL) and the number of genes inferred to be positively selected and rapidly evolving. **D.** Volcano plot showing the *d_N_*/*d_S_* rate ratio (Log_2_) and likelihood ratio test (LRT) *P*-value (–Log_10_) for each gene inferred from the best-fitting ABSREL (right) and BUSTED (left) tests in the elephant lineage. Red dots indicate genes with *d_N_*/*d_S_*>1 and *P*≤0.05, and dots in pink indicate genes with *d_N_*/*d_S_*>1 and *P*=0.051–0.20. **E.** Venn diagram showing the number and overlap of genes inferred to be positively selected (PS) or rapidly evolving (RE) by the ABSREL and BUSTED tests. **Figure 1 – source data 1. Genomes used in selection tests.** **Figure 1 – source data 2. List of positively selected genes from ABSREL.** **Figure 1 – source data 3. List of rapidly evolving genes from ABSREL.** **Figure 1 – source data 4. List of positively selected genes from BUSTED. Figure 1 – source data 5. List of rapidly evolving genes from BUSTED.** **Figure 1 – figure supplement 1. Genomic features of positively selected and rapidly evolving genes.**

Including synonymous rate variation across sites improved model fit for all genes; however, only 31 genes had significant evidence for multi-nucleotide mutations (**Figure 1C**). These data are consistent with previous observations synonymous substitution rate variation is common (Pond and Muse, 2005; Wisotsky et al., 2020) and that shorter branches tend to be more impacted by multiple hits than longer branches (Lucaci et al., 2023b, 2021) such as the elephant lineage. The best-fitting ABSREL model identified 393 genes with a class of sites with ω>1 at *P*≤0.05 in the elephant lineage (**Figure 1C**), whereas the best-fitting BUSTED model identified 427 genes with a class of sites with ω>1 at *P*≤0.05 in the elephant lineage (**Figure 1C**); 324 genes were identified as positively selected by both methods (**Figure 1D**). We also defined a set of rapidly evolving genes with a class of sites with ω>1 in either the ABSREL or BUSTED methods, but at *P*=0.051–0.20; these genes did not meet the statistical significance threshold for positive selection. However, the ABSREL and BUSTED methods are conservative, and they are likely rapidly evolving. ABSREL and BUSTED identified 127 and 608 rapidly evolving genes (**Figure 1C**), respectively, with 75 genes identified as rapidly evolving by both methods (**Figure 1D**). Positively selected and rapidly evolving genes were longer than the background gene set (**Figure 1 – Figure Supplement 1**), consistent with previous observations that ABSREL and BUSTED have more statistical power to detect ω>1 in longer genes (Murrell et al., 2015b; Pond et al., 2011b).

### Amino acid substitutions in positively selected genes are likely function-altering

To infer the putative functional consequences of amino acid changes in genes with signatures of positive selection and rapid evolution, we used IQTREE2 (Minh et al., 2020) to reconstruct the ancestral *Paenungulate* (*AncPaenungulata*) amino acid sequence for each gene (**Figure 1A**) and identified amino acid changes in the elephant lineage. Then, we used a fixed effect likelihood (FEL) model (Pond and Frost, 2005) to estimate the strength and direction of selection acting on each of these codon sites across mammals; codons with *d_N_*/*d_S_*<1 evolve under selective constraints (i.e., selection against amino acid changes). Thus, substitutions at these sites are likely function-altering. In contrast, codons with *d_N_*/*d_S_*>1 evolve under diversifying selection (i.e., selection favors amino acid changes), and codons with *d_N_*/*d_S_*≈1 evolve under weak or absent selective constraint. We found that 40.5% (4048/9981) of elephant-specific amino acid changes in positively selected genes occurred at sites with *d_N_*/*d_S_*<1, whereas 2.5% (251/9981) occurred at sites with *d_N_*/*d_S_*>1, and 47.4% (4728/9981) occurred at sites with *d_N_*/*d_S_*≈1 (**Figure 2A**). Similarly, 38.4% (7868/20366) of elephant-specific amino acid changes in rapidly evolving genes occurred at sites with *d_N_*/*d_S_*<1, whereas 2.3% (462/20366) occurred at sites with *d_N_*/*d_S_*>1, and 50.4% 10262/20366) occurred at sites with *d_N_*/*d_S_*≈1 (**Figure 2A**).

**Figure 2.**
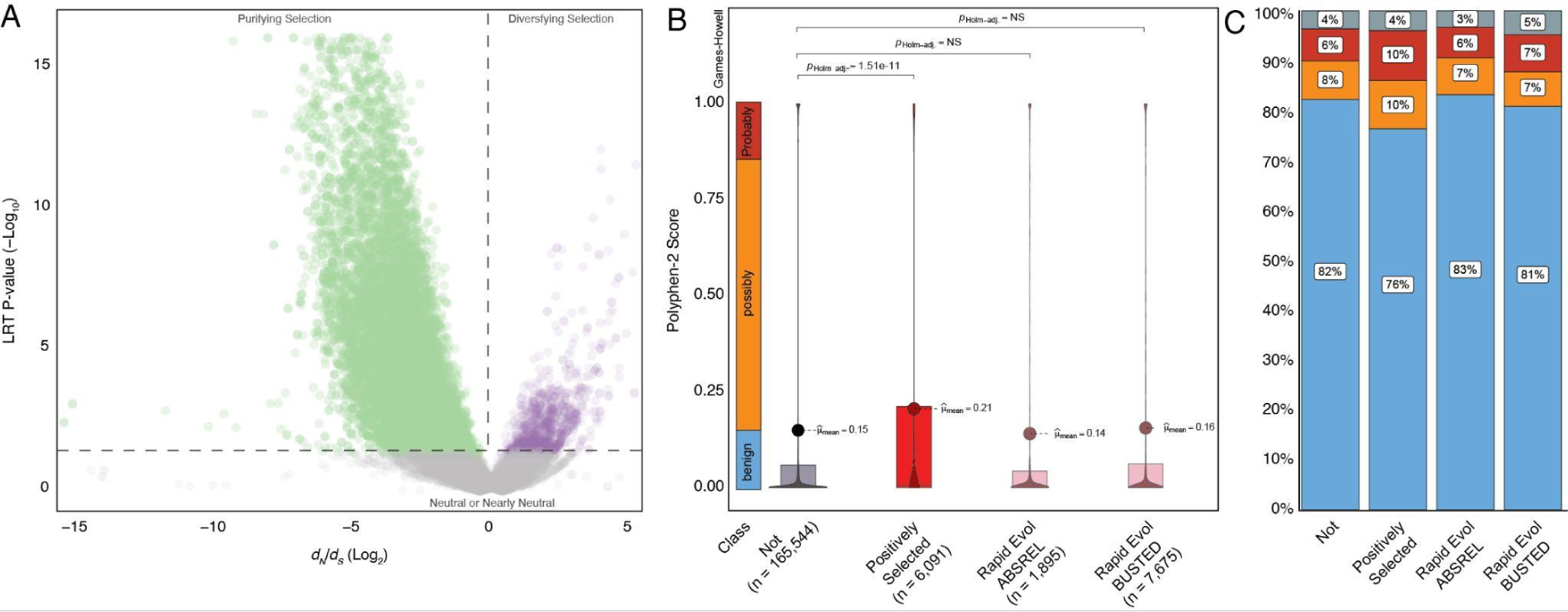
Amino acid substitutions in positively selected genes are likely deleterious. **A.** Volcano plot showing the *d_N_*/*d_S_* (Log_2_) and likelihood ratio test (LRT) *P*-value (–Log_10_) for codons with elephant-specific amino acid changes in positively selected and rapidly evolving genes. *d_N_*/*d_S_* rates and *P*-values were estimated with the fixed effect likelihood (FEL) model, the P-value is for the LRT of *d_N_*/*d_S_* ≠ 1 at each codon. Codons with *d_N_*/*d_S_*>1 and a *P*-value≤0.05 are colored purple, those with *_N_*/*d_S_*<1 and a *P*-value≤0.05 are colored green, and those with *d_N_*/*d_S_* ≈ 1 are colored grey. **B.** Violin and box plots showing the distribution of PolyPhen-2 scores for elephant-specific amino acid changes. The number of amino acid substitutions is given in parenthesis for each gene set. The range of scores corresponding to benign, possibly damaging, and probably damaging mutations is highlighted. Multiple hypotheses (Holm) corrected p values (pHolm-adj.) are shown for statistically significant comparisons; NS, not significant. Not, amino acid changes in genes with no evidence of positive selection or rapid evolution; Positively selected, amino acid changes in positively selected genes; Rapid Evol ABSREL, amino acid changes in rapidly evolving genes inferred from ABSREL; Rapid Evol BUSTED, amino acid changes in rapidly evolving genes inferred from BUSTED. **C.** Bar chart showing the qualitative PolyPhen-2 predictions of amino acid changes. Not, amino acid changes in genes with no evidence of positive selection or rapid evolution; Positively selected, amino acid changes in positively selected genes; Rapid Evol ABSREL, amino acid changes in rapidly evolving genes inferred from ABSREL; Rapid Evol BUSTED, amino acid changes in rapidly evolving genes inferred from BUSTED.

Our observation that many elephant-specific amino acid substitutions in positively selected and rapidly evolving genes occur at sites with an evolutionary signature of purifying selection (**Figure 2A**) suggests these substitutions are function-altering. To explore this possibility, we used PolyPhen-2 (Adzhubei et al., 2013) to predict the consequences of amino acid substitutions in the elephant lineage; PolyPhen-2 assigns each substitution a score in the range of 0 (benign or unlikely to be function-altering) to 1 (probably deleterious or likely to be function altering). Of 182,333 amino acid changes in the elephant lineage, 175,524 had PolyPhen-2 predictions. We combined genes identified under positive selection by ABSREL and BUSTED into a single gene set because there was considerable overlap between genes (**Figure 1C**) and found that the mean PolyPhen-2 score of amino acid changes in positively selected genes was 0.21 while the mean score of genes without evidence of selection was 0.15 (Holm adjusted Games-Howell test *P*=1.91×10^-11^); however, there was no difference in mean PolyPhen-2 scores between genes without evidence of selection and rapidly evolving genes identified by ABSREL or BUSTED (**Figure 2B**). A greater proportion of amino acid substitutions in positively selected genes are also classified as possibly and probably damaging (**Figure 2C**). These data indicate that many amino acid changes in positively selected genes occurred at sites under purifying selection and are likely function-altering.

### Positively selected and rapidly evolving genes are enriched in anti-cancer functions

Elephants have evolved cells with derived anti-cancer phenotypes; for example, they induce apoptosis at low levels of DNA damage (Abegglen et al., 2015; Sulak et al., 2016a; Vazquez et al., 2018), are resistant to oxidative stress-induced cell-death (Gomes et al., 2011), are resistant to experimental immortalization (Fukuda et al., 2016; Gomes et al., 2011), and repair DNA damage faster than smaller-bodied species (Francis et al., 1981; Hart and Setlow, 1974; Promislow, 1994). Therefore, we identified gene ontologies (GO), Reactome (Jassal et al., 2020), Wikipathway (Jassal et al., 2020), and XDeathDB (Gadepalli et al., 2021) terms related to regulated cell death modes, senescence, oxidative stress, DNA damage repair, and the cell cycle, and DisGeNET terms (Piñero et al., 2020) related to neoplasia, malignancy, and metastasis. Next, we used the hypergeometric over-representation (ORA) test in WebGestalt (Liao et al., 2019) to determine if positively selected and rapidly evolving genes were enriched in these terms and pathways. We again combined genes identified under positive selection by both methods into a single gene set and tested rapidly evolving genes identified by ABSREL or BUSTED separately. Positively selected and rapidly evolving genes were statistically enriched in numerous ontologies and pathways (FDR≤0.10) related to multiple cell death modes, regulation of the cell cycle, DNA damage repair, and cancer biology, but only one enriched term was related to oxidative stress and no enriched terms were related to senescence (**Figure 3**). Next, we tested if positively selected genes were enriched in any GO biological process or molecular function terms, i.e., not using an *a priori-defined* set of terms as above, and identified four statistically enriched GO terms at an FDR≤0.14 (**Figure 4**). Similarly, rapidly evolving genes identified by ABSREL were enriched in numerous GO terms at an FDR≤0.10 (**Figure 4**), most of which were related to immune functions. In contrast, rapidly evolving genes identified by BUSTED were not enriched in any ontology term with an FDR≤0.94. Thus, we conclude that genes related to cell death modes, cancer biology, oxidative stress, senescence, EGFR-signaling, and immune functions, particularly neutrophil biology, have been pervasive targets of positive selection and rapid evolution in the elephant lineage.

**Figure 3.**
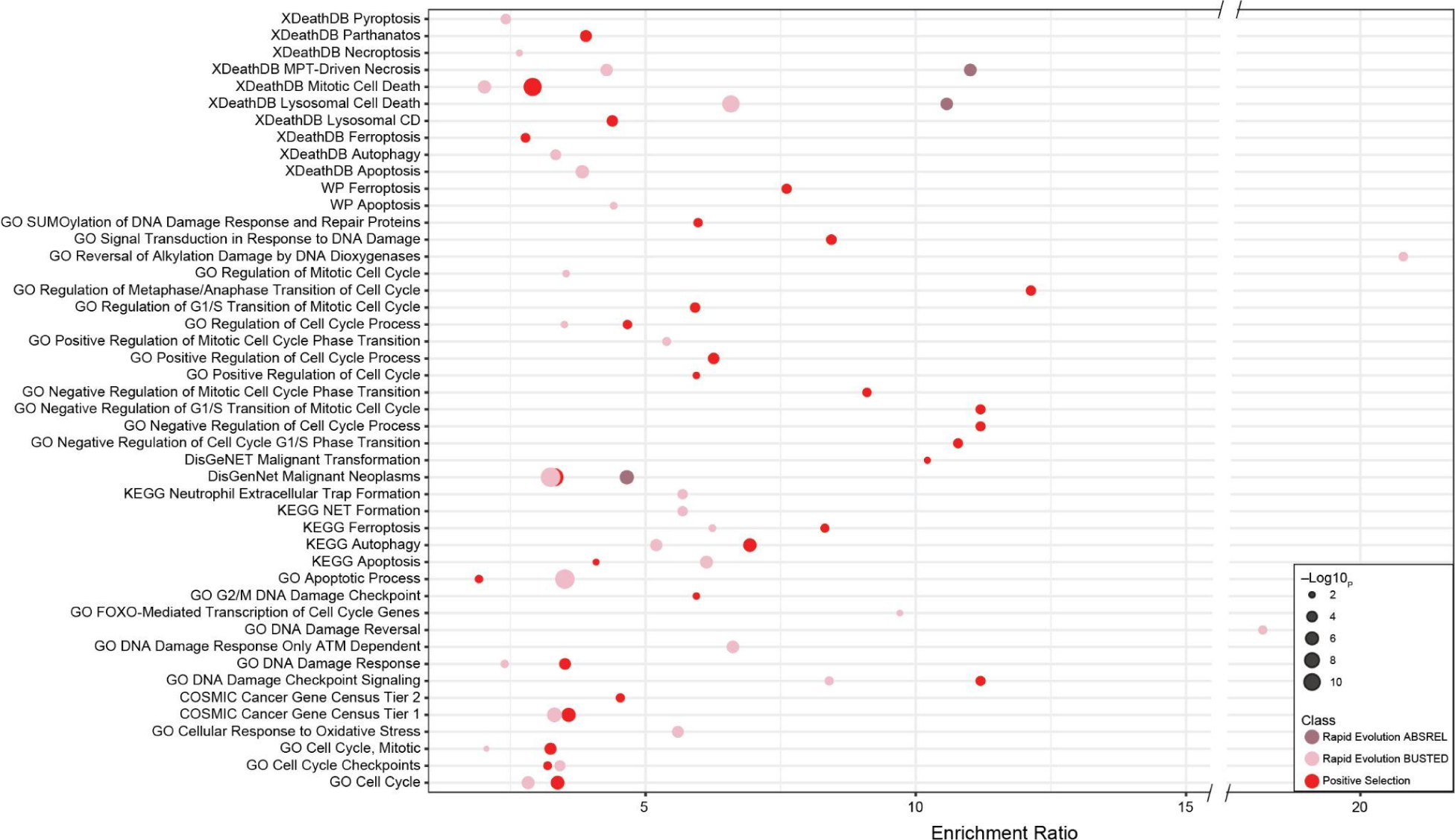
Positively selected and rapidly evolving genes are enriched in cell death pathways and cancer gene sets. Bubble plot showing the enrichment ratio and hypergeometric *P*-value of GO, Reactome (R), Wikipathway (WP), and XDeathDB terms related to regulated cell death modes, senescence, oxidative stress, DNA damage repair, and the cell cycle, and DisGeNET terms related to neoplasia, malignancy, and metastasis in which positively selected (red) and rapidly evolving (pink) genes are enriched (FDR≤0.10). Figure 3 **– source data 1**. Webgestalt output files for positively selected genes. Figure 3 **– source data 2**. Webgestalt output files for ABSREL rapidly evolving genes. Figure 3 **– source data 3**. Webgestalt output files for BUSTED rapidly evolving genes.

**Figure 4.**
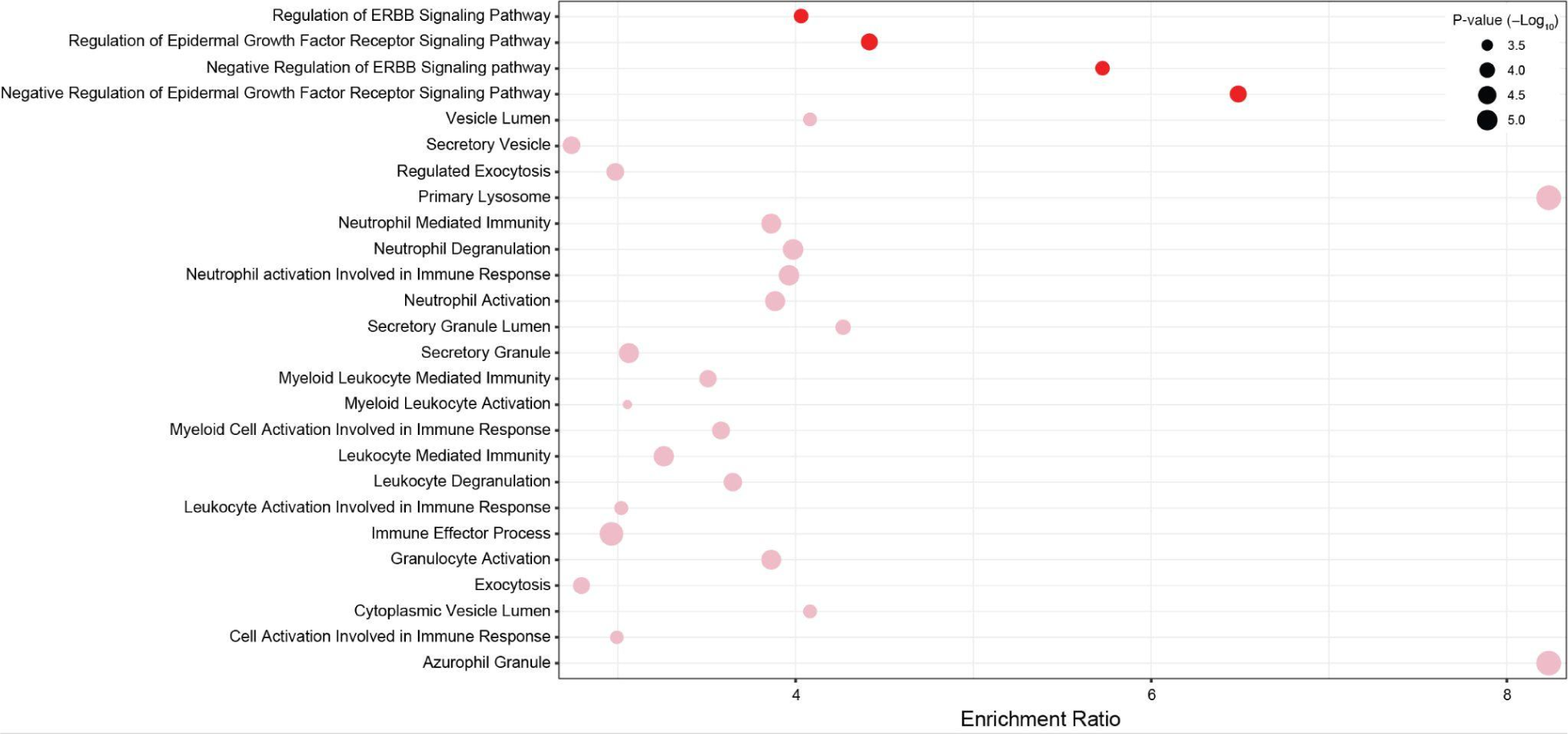
Gene ontology (GO) terms in which positively selected and rapidly evolving genes are enriched. Bubble plot showing the GO biological process and molecular function name, enrichment ratio, and hypergeometric P-value of terms in which positively selected (red, FDR≤0.14) and rapidly evolving identified by ABSREL (pink, FDR≤0.10) genes are enriched. Figure 4 **– source data 1**. Webgestalt output files for positively selected genes. Figure 4 **– source data 2**. Webgestalt output files for ABSREL rapidly evolving genes. Figure 4 **– source data 3**. Webgestalt output files for BUSTED rapidly evolving genes.

## Discussion

Elephants have a very low prevalence of cancer given their body size and lifespan (Abegglen et al., 2015; Bulls et al., 2022; Compton et al., 2023; Tollis et al., 2021), indicating that elephants have evolved enhanced cancer prevention mechanisms. Among these anti–cancer phenotypes are cells that induce apoptosis at low levels of DNA damage (Abegglen et al., 2015; Sulak et al., 2016a; Vazquez et al., 2018), are resistant to oxidative stress-induced cell death (Gomes et al., 2011), have faster DNA damage repair rates than smaller-bodied species (Francis et al., 1981; Hart and Setlow, 1974; Promislow, 1994), and resistant to experimental immortalization (Fukuda et al., 2016; Gomes et al., 2011). These cellular traits are at least partly mediated by an increase in the number of tumor suppressors in the elephant lineage (Caulin et al., 2015; Doherty and Magalhães, 2016; Sulak et al., 2016b; Tollis et al., 2020; Vazquez et al., 2018; Vazquez and Lynch, 2021), but many other mechanisms must also have contributed to enhanced cancer resistance including functional divergence of protein-coding genes (Li et al., 2023; Lynch and Wagner, 2008). Here, we used a suite of methods to characterize the strength and direction of selection acting on protein-coding genes in the elephant lineage, focusing on genes with evidence of positive selection and rapid evolution, which is likely associated with functional divergence (Slodkowicz and Goldman, 2020; Tennessen, 2008). We identified 496 genes, 3.73% of alignments tested, with statistically significant evidence for positive selection in the elephant lineage, and 660 genes, 4.96% of alignments tested, that likely evolved rapidly in elephants. These are good candidates for genes that may contribute to elephant-specific traits, such as the evolution of molecular and cellular characteristics that enhance cancer resistance, among many others.

### Comparison to previous studies of selection in elephants

Several previous studies have explored the role of positive selection acting on protein-coding genes in the elephant lineage, albeit with smaller numbers of species and alignments than our study or using different methods. Goodman et al. (2009), for example, used the free ratio model in PAML (Yang, 2007) to identify positively selected genes in the elephant lineage using a dataset of 11 species and 5,714 protein-coding genes; they identified 67 genes with *d_N_*/*d_S_*≥1.01, which were enriched in mitochondrial functions but none of the GO terms we identified (**Supplementary dataset 1**). There are several important differences between Goodman et al. (2009) and our study: 1) The free ratio model does not test if the *d_N_*/*d_S_* ratio in a particular lineage is significantly different than one, rather it tests if *d_N_*/*d_S_* rate variation is different across lineages (Yang, 1998); 2) Unlike ABSREL and BUSTED, the free ratio model does not allow for rate variation across sites, *d_N_*/*d_S_*>1 is only inferred if the average *d_N_*/*d_S_* across all sites is greater than 1; and 3) The method implemented in PAML does not account for synonymous rate variation across sites or multiple simultaneous codon substitutions, which can bias inferences of both *d_S_ (Dunn et al., 2019; Rahman et al., 2021; Wisotsky et al., 2020)* and *d_N_* (Dunn et al., 2019; Lucaci et al., 2023a; Nozawa et al., 2009; Suzuki, 2008; Venkat et al., 2018; Yang and Reis, 2011). While we tested 81.6% (4,663) of the genes tested by Goodman et al. (2009), only one (*CD3G*) was inferred to have been positively selected in our analyses; therefore, it is unsurprising that there was little overlap in our GO enrichment results.

We previously reported the results of a preliminary selection scan using the same dataset of alignments used here and the base ABSREL model, which did not include synonymous rate variation or multiple hits (Raviv et al., 2023). We identified 674 positively selected genes enriched in several Panther pathways (Mi et al., 2013; Mi and Thomas, 2009; Thomas et al., 2022) related to neurotransmitter signaling (Raviv et al., 2023). However, we found in this study that synonymous rate variation across sites improved model fit for all genes, suggesting that at least some of the 674 genes we previously reported as positively selected may be false positives; indeed, only 215 (31.9%) were inferred as positively selected by ABSREL[S]. However, the 393 genes identified as positively selected by ABSREL[SMH] were similarly enriched in Panther pathways related to neurotransmitter signaling (**Supplementary dataset 2**), particularly serotonin and muscarinic acetylcholine receptor signaling. Thus, the signal for positive selection on genes in these pathways is strong despite the small overlap between genes inferred as positively selected by ABSREL and ABSREL[S].

Li et al. (2023) used BUSTED[S] to test for an episode of positive selection in the stem lineage of African and Asian elephants in a dataset of 12 species and 4,444 protein-coding genes and identified 618 positively selected genes. 492 of these genes were also tested for selection in our dataset, but only 32 (6.45%) were inferred to have been positively selected in our analyses. Not surprisingly, given the low overlap, our datasets had no overlap in GO term or pathway enrichments (**Supplementary dataset 3**). There are several potential explanations for this low overlap, including differences in our alignments, which can have substantial impacts on inferences of selection (Fletcher and Yang, 2010; Franco et al., 2019; Jordan and Goldman, 2012; Markova-Raina and Petrov, 2011; Spielman et al., 2014) and differences in the number of species included in alignments (261 vs 12), with larger alignments likely having more power and reduced false positive inferences of selection (Pond and Frost, 2005) but also paradoxical effects on inferences of selection, for example, by diluting the signal of *d_N_*/*d_S_*>1 by the addition of taxa in which positive selection did not occur (Chen and Sun, 2011; Murrell et al., 2012; Smith et al., 2015; Yokoyama et al., 2008); this signal loss may be more likely to occur in cases where the strength of selection is relatively weak or the proportion of sites or lineages under selection relatively small (Yang and Reis, 2011). Thus, we interpret our results as complementary to, rather than in contrast with, Li et al. (2023). Indeed, studies of individual genes should include taxa inclusion sensitivity analyses (such as jackknifing or a priori exclusion of focal taxa) to infer the effects of alignment size and composition on inferences of positive selection.

### Positive selection and rapid evolution of cancer genes in elephants

We reasoned that genes with roles in cancer biology might have diverged in function in elephants, for example, coincident with the evolution of enhanced tumor suppressor or reduced oncogenic functions. We found that positively selected and rapidly evolving genes were statistically enriched in many GO terms and pathways related to cancer biology, including malignant neoplasms, neoplastic cell transformation, and COSMIC Cancer Gene Census Tier 1 and 2; Tier 1 genes have a documented activity related to cancer and evidence of mutations that promote oncogenic transformation, while Tier 2 genes genes have a strong indication of a role in cancer but with less extensive evidence than Tier 1 genes. Among these genes are many with well-characterized roles in cancer biology, such as *APC*, a negative regulator of canonical WNT signaling that suppresses invasion and tumor progression, particularly in colorectal cancer (Fodde, 2002; Hankey et al., 2018; Kwong and Dove, 2009), *CHEK1* which encodes the checkpoint kinase CHK1 that regulates multiple cell cycle and DNA damage checkpoints (Chen and Sanchez, 2004; Harten et al., 2019; Liu et al., 2000; Patil et al., 2013; Synnes et al., 2002), and *POLD1*, which encodes the catalytic subunit of the DNA polymerase δ complex that has both DNA polymerizing and a proofreading domain that ensures replication accuracy and genomic stability (Haradhvala et al., 2018); germline mutations in the proofreading domain of POLD1 predispose to colorectal adenomas and carcinomas among others (Palles et al., 2013; Rayner et al., 2016). These data suggest that functional divergence of genes with cancer-protective effects may have contributed to the evolution of cancer resistance in elephants and underlie cellular traits like rapid DNA damage repair.

### Positive selection and rapid evolution of cell death pathway genes in elephants

Many mechanisms have likely contributed to the evolution of reduced cancer risk in elephants, including cells that induce apoptosis at low levels of DNA damage (Abegglen et al., 2015; Sulak et al., 2016a; Vazquez et al., 2018) and are resistant to oxidative stress-induced cell death (Gomes et al., 2011). Consistent with these functional studies, we found that positively selected and rapidly evolving genes were statistically enriched in many GO terms and pathways related to cell death, such as apoptosis, ferroptosis, pyroptosis, and parthanatos, and regulated across (**Figure 3**). These data suggest that positively selected and rapidly evolving genes have contributed to the divergence of cell death pathways in elephants. Among the rapidly evolving genes, for example, is *CASP8*, the initiator caspase of extrinsic apoptosis, necroptosis, and pyroptosis (Boldin et al., 1996; Fritsch et al., 2019; Muzio et al., 1996), *APAF1*, which forms the central scaffold of the apoptosome and and thus plays a critical role in apoptosis (Bratton and Salvesen, 2010; Cain et al., 2002), *BOK* an unusual BCL-2 family protein that directly induces mitochondrial outer membrane permeabilization (MOMP) and apoptosis in response to endoplasmic reticulum (ER) stress (Carpio et al., 2015; Llambi et al., 2016), *BCL2L15*, another unusual BCL-2 family protein with pro-apoptotic functions that reduces malignant transformation of cells in the gastrointestinal tract (Dempsey et al., 2005; Jang et al., 2022; Özören et al., 2009)*, BCLAP* a tumor suppressor that induces cell cycle arrest at the G_1_/S and apoptosis through an uncharacterized but p53-independent mechanisms (Fan et al., 2011; Han et al., 2022; Yao et al., 2007; ZHAO et al., 2016), *KDM6B*, which promotes parthanatos by inhibiting DNA damage-evoked PARP-1 hyperactivation (Yang et al., 2022), and *ALKBH7* is required for alkylation and oxidation-induced programmed necrosis (Fu et al., 2013; Jordan et al., 2017; Kulkarni et al., 2020). Remarkably, other cancer-resistant species have also evolved modified cell death pathways. Blind mole rats, for example, evolved a concerted necrotic cell death mechanism (Gorbunova et al., 2012), and naked mole rats evolved cells that induce cell death in response to senescence via the INK4a-RB pathway (Kawamura et al., 2023). These data suggest functional divergence of cell death modes may be shared across cancer-resistant species.

### Positive selection and rapid evolution of cell cycle regulatory genes in elephants

The cell cycle and its checkpoints play crucial roles in the development and progression of cancer; loss of checkpoint control, for example, allows cells to proliferate despite DNA damage contributing to genomic instability and resistance to apoptosis (Matthews et al., 2022) Numerous genes with roles in the cell cycle were positively selected or are evolving rapidly in elephants, including *PPP2CB*, which encodes the catalytic subunit of the master regulator of the cell cycle phosphatase 2A (PP2A) (Wlodarchak and Xing, 2016), *CDKAP2*, which inhibits cell cycle progression at the G_1_/S checkpoint by suppressing CDK2 and blocking the interaction between CDK2 and cyclin E and A (Liu et al., 2013), E2F7, which represses the expression of G_1_/S genes during S-phase progression (Westendorp et al., 2012) and *CDC25B*, a phosphatase required to activate cyclin/CDK1 complexes at the G_2_/M checkpoint and induce mitotic progression (Schmidt et al., 2007). Many genes related to the centrosome, which is required for several cell-cycle transitions, including G_1_ to S-phase, G_1_ to M, and metaphase to anaphase (Doxsey et al., 2005), were positively selected or evolved rapidly, including *CEP57*, *CEP131*, *CEP76*, *CEP350*, *CEP76*, and *CEP104*. These data suggest that the functional divergence of genes that regulate the cell cycle has likely contributed to the evolution of cancer resistance in elephants.

### Positive selection and rapid evolution of DNA damage repair genes in elephants

DNA damage checkpoints are essential to maintain genomic integrity. Perhaps surprisingly, given their critical role, DNA damage responses can dramatically diverge between species. Previous studies have also found that elephant cells repair DNA damage much faster than other smaller-bodied mammals (Francis et al., 1981; Hart and Setlow, 1974; Promislow, 1994), suggesting that DNA damage repair genes may have derived functions in elephants. Among the positively selected and rapidly evolving genes, for example, are *RIF1*, which cooperates with TP53BP1 to promote DNA repair by non-homologous end joining (NHEJ) in the G_1_ and S phases of the cell cycle (Escribano-Díaz et al., 2013; Feng et al., 2013; Virgilio et al., 2013), the multifunctional exonuclease *EXO1*, which is involved in DNA damage checkpoint progression, mismatch repair (MMR), translesion DNA synthesis (TLS), nucleotide excision repair (NER), and limits end resection of double-strand breaks thereby facilitating repair via error-free homologous recombination (HR) rather than error-prone NHEJ (Keijzers et al., 2018; Tomimatsu et al., 2017), *MLH1*, part of PMS2 MMR complex that generates single-strand breaks near the mismatch and entry points for EXO1 to degrade the strand containing the mismatch (Kadyrov et al., 2006; Kansikas et al., 2011; Sacho et al., 2008), *NEK4*, which regulates a unique ATM/ATR-independent DNA damage checkpoint and the induction of replicative senescence in response to double-stranded (DSB) DNA damage (Chen et al., 2011; Tomimatsu et al., 2017), *KAT5*, an acetyltransferase that plays an essential role in DNA damage repair by acetylating and activating ATM and the canonical DSB repair pathway (Sun et al., 2005)and *SAMHD1* which promotes DNA end resection to facilitate DSB repair by HR (Daddacha et al., 2017; Kapoor-Vazirani et al., 2022). Similarly, long-lived and cancer-resistant Bowhead whales, which have a maximum lifespan of over 200 years (George et al., 1999), have evolved cells that repair double-strand breaks with high efficiency and accuracy compared to many other mammals (Firsanov et al., 2023), suggesting that the evolution of DNA damage repair genes may be a common route to evolve cancer resistance.

### Positive selection of EGFR-signaling genes in elephants

While this study was motivated by the evolution of cancer resistance in the elephant lineage, our genome-wide approach is relatively unbiased with respect to the role of protein-coding genes in cancer, i.e., we tested for selection in all genes rather than genes with previously ascertained roles in cancer biology. Thus, our observation that positively selected genes were only enriched in four GO biological process terms (with an FDR<0.10), and that after redundancy reduction only “regulation of epidermal growth factor receptor signaling pathway” remained is remarkable. Dysregulated epidermal growth factor receptor (EGFR) signaling plays a central role in many cancers, including driving tumor growth and metastasis (Normanno et al., 2006; Sigismund et al., 2018), and EGFR inhibitors are a standard treatment for cancers with oncogenic EGFR mutations (Cooper et al., 2022; Jänne et al., 2015; Levantini et al., 2022). Among the positively selected genes in the EGFR pathway identified by ABSREL and BUSTED is *SPRY2*, a negative regulator of receptor tyrosine kinases (RTKs) and the Ras/Erk signaling pathway (Guy et al., 2009; Hacohen et al., 1998), that functions as a context-dependent tumor suppressor or oncogene (Holgren et al., 2010; Lo et al., 2006; Saini et al., 2018). For example, epigenetic silencing of *SPRY2* is common in prostate cancer (McKie et al., 2005), and it is inactivated in about 18% of primary and over 70% of metastatic tumors (Taylor et al., 2010); SPRY2 cooperates with PTEN and PP2A to act as a tumor suppressor checkpoint in prostate cancer (Patel et al., 2013). In contrast, *SPRY2* expression is upregulated in colonic adenocarcinomas, where it promotes cell proliferation, growth of tumor xenografts, metastasis, angiogenesis, and inhibits apoptosis (Holgren et al., 2010). These data suggest a relatively direct connection between positively selected genes in the EGFR–signaling pathway and the evolution of reduced cancer prevalence in elephants.

### Derived neutrophil biology in elephants?

Our observation that positively selected and rapidly evolving genes in elephants are enriched in gene ontologies related to neutrophil biology is particularly intriguing. While neutrophils are best known for their role in the innate immune system, they also play an important, if complex, role in cancer biology, where they can both promote and inhibit tumor initiation, growth, and metastasis (Coffelt et al., 2016). Remarkably, many positively selected and rapidly evolving genes are related to neutrophil extracellular trap formation, a form of cell death characterized by the release of decondensed chromatin and granule contents into the extracellular space (Poli and Zanoni, 2023; Remijsen et al., 2011; Vorobjeva and Chernyak, 2020), granules and degranulation, and neutrophil granule proteins have been associated with tumor progression (Mollinedo, 2019). Neutrophils contain (at least) four functionally distinct kinds of granules: 1) primary or azurophilic granules, which store proteins with antimicrobial activity such as myeloperoxidase, elastase, cathepsins, and defensins; 2) secondary or specific granules, which contain lactoferrin and lipocalin, but not matrix metalloprotease-9 (MMP9); 3) tertiary granules, which contain MMP9 but not lactoferrin or lipocalin; and 4) secretory vesicles, which are likely derived from endocytosis of the plasma membrane (Lacy, 2006).

We found that rapidly evolving genes were enriched in GO cellular component terms related to each of these kinds of granules (**Figure 4**), including granule contents such as *CTSS*, *LYZ*, *MMP9*, and *PRDX6*, which might be expected to evolve rapidly because they are caught in an arms race with pathogens, but also genes that mediate degranulation including members of the SNAP receptor (SNARE) signaling pathway *SNAP23*, *VAMP2*, *VAMP7*, *VAMP8*, *GCA*, *STX5*, and *STXBP2* which might be expected to have conserved functions given their essential roles in conserved biological processes such as membrane trafficking. These data suggest neutrophil phenotypes related to cancer biology may have evolved in the elephant lineage. Indeed, some data indicate that neutrophil biology is different in elephants than in other species. Elephants, for example, have higher concentrations of neutrophils in their blood than expected from their body mass, i.e., neutrophil concentration in blood scales hypermetrically with body size (Downs et al., 2019). Elephant neutrophils are also activated and produce oxygen radicals at much lower stimuli than other species (Smith et al., 1998; Tell et al., 1999). Further studies are needed, however, to determine if these phenotypes are derived in the elephant lineage and to explore if other neutrophil functions such as neutrophil extracellular trap formation (NETosis), a regulated cell death mode characterized by intracellular degranulation and the release of decondensed chromatin and granular contents into the extracellular space (Metzler et al., 2014, 2011; Papayannopoulos et al., 2010; Vorobjeva and Chernyak, 2020), are different in elephants than other species.

### Caveats and limitations

There are several limitations to this study. For example, while there are two major lineages of living elephants, African (*Loxodonta africana* and *Loxodonta cyclotis*) and Asian (*Elephas maximus* ssp.), our alignments only included African savanna elephant (*Loxodonta africana*). Therefore, our inference of selection combines codon substitutions in the *Proboscidean* stem lineage with those in the *Loxodonta* lineage after the *Loxodonta*–*Elephas* divergence, which cannot have contributed to the evolution of elephant-specific traits. We do not believe that this adversely affects our results because the length of the Proboscidean stem lineage (∼53 million years) is much longer than the divergence between the African and Asian elephant lineages, which is estimated to be 6.6–8.6 MYA (Rohland et al., 2007). Thus, most amino acid changes will have occurred in the *Proboscidean* stem lineage; codon-based models also have little power to detect recent selection. Perhaps most importantly, the ability of codon-based methods to detect positive selection is sensitive to a range of parameters, including branch length (substitutions per codon), synonymous site saturation, the strength of selection, and the number of selected sites. Tests of positive selection, for example, are conservative and lack power when the strength of selection is weak, limited to one or a few sites, and episodic, i.e., occurs on a single lineage (such as the *Proboscidean* stem lineage). In these cases, the methods lack the power to reject the null model, and an episode of positive selection will be missed. Thus, our results are not an exhaustive catalog of all genes positively selected in elephants. Future studies could use other methods to detect the signature of protein functional divergence and episodic purifying selection (Gaucher et al., 2002; Gu, 2006, 2001a, 2001b; Gu et al., 2013; Philippe et al., 2003; Pond et al., 2020); these methods, however, would need to include many more *Proboscidean* species, including living species such as Asian and African forest elephants and extinct species with genomic data such as straight-tusked elephants, mammoths, and mastodons, to have power to detect shifts in amino acid conservation profiles.

### Conclusions

Elephants have emerged as a model system to study the evolution of cancer resistance in large and/or long-lived species. While numerous molecular and cellular mechanisms must have contributed to cancer resistance and other traits in the elephant lineage, in this study, we focused on identifying genes that were positively selected or evolved rapidly and may have functionally diverged in elephants. Our results indicate that genes related to many cell death pathways, EGFR–signaling, and neutrophil biology were positively selected or evolved rapidly in the elephant lineage. At least some of the genes likely contributed to the evolution of cancer resistance in *Proboscideans* and are promising candidates for functional studies. More generally, our results highlight the importance of including variation in synonymous substitution rates and multi-nucleotide mutations in *d_N_*/*d_S_*-based selection tests; indeed, we found statistically significant evidence for synonymous substitution rate variation in all genes we tested.

## Methods

### Assembling alignments of orthologous coding genes from 261 Eutherians

We previously reported the assembly of protein-coding gene alignments from the genomes of 261 *Eutherian* (“Placental”) mammals (Bowman et al., 2023b), which are also used in this study. Briefly, we used a reciprocal best BLAT hit (RBBH) approach to assemble a dataset of orthologous coding gene alignments from the genomes of 261 *Eutherian* (“Placental”) mammals (**Figure 1A**; **Figure 1 – source data 1**). We began by using RBBH with query sequences from human CDSs in the Ensembl v99 human coding sequence dataset and BLAT searching the genomes of 260 other Eutherians, followed by searching the top hit to the human query back to the human genome; we used BLAT matching all possible reading frames, with a minimum identity set at 30% and the “fine” option activated (Kent, 2002), and excluded genes with fewer than 251 best reciprocal hits out of the 261 (human+other mammals) species included in the analysis. Orthologous genes were aligned with Macse v2 (Ranwez et al., 2018), a codon-aware multiple sequence aligner that identifies frameshifts and readjusts reading frames accordingly. Alignments generated by Macse v2 were edited by HMMcleaner with default parameters (Franco et al., 2019) to remove species-specific substitutions that are likely genome sequencing errors and “false exons’’ that might have been introduced during the Blat search. Finally, we excluded incomplete codons and the flanks of indels, which usually have more misaligned substitutions. We thus generated a dataset of 13,310 orthologous coding gene alignments from the genomes of 261 *Eutherian* mammals; this corresponds to 66.4% of all protein-coding genes annotated in the African savannah elephant genome (Loxafr3.0).

### ABSREL and BUSTED selection tests

The multiple sequence alignments used in this study have been described elsewhere (Bowman et al., 2023b). Methods to infer the strength and direction of selection acting on molecular sequences, such as those to identify pervasive and episodic adaptive evolution (“positive selection”), are often based on estimating the ratio of non-synonymous (*d_N_*) to synonymous (*d_S_*) substitution rates (*d_N_*/*d_S_* or *ω*) in an alignment of homologous genes. These methods can be applied to entire genes or regions within genes (Hughes and Nei, 1988), sites (Nielsen and Yang, 1998), branches (Messier and Stewart, 1997; Muse and Gaut, 1994; Yang, 1998), or sites along a specific branch - the latter often called “branch-site” (BSREL) models (Pond et al., 2011a; Smith et al., 2015; Yang and Nielsen, 2002). An advantage of the BSREL models is that they allow for a group of sites to evolve under the action of positive selection, i.e., with *ω*>1, while the remaining sites can evolve under purifying selection (*ω*<1) or relatively neutrally (*ω*=1) in specific branches, and thus can detect episodic positive selection (limited to a subset of branches) and restricted to a subset of sites in a gene. Positive selection is inferred when a class of sites is identified with *ω*>1, with a LRT P-value ≤ 0.05. We used two related BSREL models implemented in HyPhy (Pond et al., 2005), ABSREL, which infers the number of *d_N_*/*d_S_* rate categories to be inferred from the alignment, and BUSTED, which has three fixed rate classes. ABSREL and BUSTED can accommodate both synonymous rate variation across sites [S] (Pond and Muse, 2005; Wisotsky et al., 2020) and multi-nucleotide mutations per codon [MH] (Lucaci et al., 2021). We tested for selection using the base ABSREL and BUSTED models, models that account for synonymous rate variation across sites (ABSREL[S] and BUSTED[S]), models that account for multi-nucleotide mutations per codon (ABSREL[MH] and BUSTED[MH]), and models that account for both synonymous rate variation across sites and multi-nucleotide mutations per codon (ABSREL[SMH] and BUSTED[SMH]). HyPhy v2.5.36 was used to run ABSREL v2.3 and BUSTED v3.1, while HyPhy v.5.32 was used to run FEL v2.1. Positive selection is inferred when the best-fitting ABSREL or BUSTED model, determined from a small sample AIC test, includes a class of sites with *ω*>1 at a likelihood ratio test (LRT) *P*-value ≤ 0.05, and rapid evolution inferred when the best-fitting ABSREL or BUSTED models inferred a class of sites with *ω*>1 at a likelihood ratio test (LRT) *P*-value = 0.051 – 0.20.

### Fixed Effects Likelihood (FEL) selection tests

We characterized the strength and direction of selection acting on each codon site in positively selected and rapidly evolving genes in elephant lineage with a fixed effects likelihood model (FEL) that included synonymous rate variation across sites (Pond and Frost, 2005). This model estimates separate *d_N_*/*d_S_* rates for each site in an alignment and quantifies the strength of selection. We used these per site *d_N_*/*d_S_* rates to estimate the likely functional consequences of amino acid substitutions at those sites, such that substitutions at sites with *d_N_*/*d_S_* < 1 are likely to be function-altering and deleterious, substitutions at sites with *d_N_*/*d_S_* = 1 are likely to be non-function altering, and substitutions at sites with *d_N_*/*d_S_* > 1 are likely to be function altering and adaptive.

### Identification of elephant-specific amino acid changes

We used IQTREE2 (Nguyen et al., 2015) to reconstruct ancestral amino acid sequences of all 13,491 genes to identify elephant-specific amino acid changes. For each gene, we translated nucleotide sequences to amino acid sequences with the -st NT2AA option, imposed the species tree with the *Paenungulate* lineages as polytomy with the -te option, inferred the best model of amino acid substitution and rate variation across sites with the -m MFP option (Kalyaanamoorthy et al., 2017), and to infer ancestral protein sequences with the --ancestral option; identical protein sequences were maintained in the analyses with the --keep-ident option to ensure consistent internal node numbering across genes. After the reconstructions for each gene were completed, the predicted ancestral *Paenungulate* (*AncPaenungulata*), i.e., the last common ancestor of elephant, manatee, and hyraxes, of each gene was compared to the *Loxodonta africana* sequence to identify amino acid changes in *Loxodonta africana*. We thus identified 182,333 amino acid changes in 13,310 proteins in the elephant lineage.

### PolyPhen-2 amino acid classification

We used PolyPhen-2 (Adzhubei et al., 2013) to classify the functional consequences of amino acid changes in the gorilla lineage using the *AncPaenungulata* (**Figure 1A**) reconstructed protein sequences as input and “wildtype” amino acid and amino acid changes in elephant lineage as the “mutant” amino acid. The Classifier model was set to “HumDiv”, and the other parameters to the GRCh37/hg19 genome, canonical transcripts, and missense annotations.

### Over Representation Analyses (ORA)

We combined genes identified as positively selected by ABSREL (n=393) and BUSTED (n=427) into a single gene set because 82% (324/393) of genes identified as positively selected by ABSREL were also identified as positively selected by BUSTED, whereas 76% (324/427) of genes identified as positively selected by BUSTED were also identified as positively selected by ABSREL. In contrast, only 59% (75/127) and 12% (75/608) of genes were classified as rapidly evolving by both ABSREL and BUSTED, respectively. Thus, we analyzed these gene sets independently.

We identified gene ontologies (GO), Reactome (Jassal et al., 2020), Wikipathway (Jassal et al., 2020), and XDeathDB (Gadepalli et al., 2021) terms related to regulated cell death modes, senescence, oxidative stress, DNA damage repair, and the cell cycle, and DisGeNET terms (Piñero et al., 2020) related to neoplasia, malignancy, and metastasis directly from each database or with the term search function in Enrichr (Chen et al., 2013; Kuleshov et al., 2016). We used the hypergeometric over-representation (ORA) test in WebGestalt (Liao et al., 2019) to determine if positively selected and rapidly evolving genes were enriched in these terms and pathways. We combined genes identified under positive selection by both methods into a single gene set, tested rapidly evolving genes identified from ABSREL and BUSTED independently, the minimum number of genes for a category was set to 5, we used as the background gene set all genes tested for selection, and controlled false discovery with the Benjamini-Hochberg false discovery rate (Benjamini and Hochberg, 1995).

We also used WebGestalt (Liao et al., 2019) to identify if positively selected and rapidly evolving genes were enriched in any gene ontology (GO) Biological Process and Cellular Component ontologies without a priori selecting candidate gene sets. Because many genes were identified as positively selected or rapidly evolving, we set the minimum number of genes for a category to 30 for tests without a priori identified GO terms and pathways, which excludes GO terms with fewer than 30 annotated genes; this strategy reduces the likelihood of inferring enrichment for ontologies with few annotated genes by chance because our foreground gene sets are large.

### Characteristics of positively selected and rapidly evolving genes

We used ShinyGO 0.77 (Ge et al., 2019) to characterize the high-level functions (defined by GO terms) and genomic features of positively selected and rapidly evolving genes; as for the ORA described above, we analyzed positively selected genes as a single group but rapidly evolving gene sets independently. Positively selected and rapidly evolving genes were longer than the background gene set (**Figure 1 – figure supplement 1**), consistent with previous observations that ABSREL and BUSTED have more statistical power to detect deviation from ω>1 in longer genes (Murrell et al., 2015b; Pond et al., 2011b), positively selected genes were also more GC rich than the background gene set (**Figure 1 – figure supplement 1**).

### Comparison to previous studies of selection in elephants

#### Comparison to Goodman et al. (2009)

Goodman et al. (2009) used the free ratio model implemented in the codeml program of PAML (Yang, 1997a, 1997b) to identify positively selected genes in the elephant lineage using a dataset of 11 species and 7,768 RefSeq transcripts corresponding to ∼6,000 nonredundant protein-coding genes in 2009; note that this is more properly a test of *d_N_*/*d_S_* rate variation across lineages, and not a statistical test of positive selection because the hypothesis of selection is not directly tested against the null model of *d_N_*/*d_S_*=1; it is also unclear if the free ratio model was statistically a better fit to the data than a simple one-ratio model, which is the appropriate null model against which the free ratio model is usually tested (Yang, 1998). However, the free ratio model is a branch-based test that does not account for rate variation across sites and is, therefore, a very conservative test for positive selection.

To ensure a fair comparison between our results, particularly our GO term enrichment results, we downloaded their codeml results from Dryad (https://doi.org/10.5061/dryad.908) and used BioMart (https://www.ensembl.org/info/data/biomart/index.html) to convert ReSeq IDs to HUGO gene IDs based on Ensembl Genes 111 and GRCh38.p14; the redundancy reduced dataset included 5,714 protein-coding genes, of which 67 had *d_N_*/*d_S_*≥1.01. We again used WebGestalt v. 2019 (Liao et al., 2019) to identify if genes with *d_N_*/*d_S_*≥1.01 were enriched gene ontology (GO) Biological Process and Cellular Component ontologies using the over-representation analysis (ORA) with the minimum number of genes for a category set to 5 (default) 30 (as in our analyses), the 67 genes with *d_N_*/*d_S_*≥1.01 as the foreground and the 5,714 genes tested for selection as the background gene set.

The 67 rapidly evolving genes (i.e., those with *d_N_*/*d_S_*≥1.01) were reported to be enriched in mitochondrial functions (Goodman et al., n.d.), which is similar to previous studies that have reported rapid evolution of mitochondrially encoded protein-coding genes in elephants (Finch et al., 2014; Ngatia et al., 2019), but not in any of the GO terms we identified (Goodman et al., n.d.). Goodman et al. (2009) used an early version of DAVID (Sherman et al., 2007) to identify enriched GO terms, suggesting that differences in our results may arise from differences in our enrichment tests. Therefore, we used WebGestalt to reanalyze GO term enrichment for the 67 rapidly evolving genes identified by Goodman et al. (2009) and found they were enriched (hypergeometric *P*≤0.05) in many GO biological process and cellular component terms related to the mitochondria (**Supplementary dataset 1**), but none the terms we identified.

#### *Comparison to* Li et al. (2023)

Li et al. (2023) used BUSTED[S] to test for an episode of positive selection in the stem lineage of African and Asian elephants in a dataset of 12 species and 4,444 protein-coding genes and identified 618 positively selected genes. To ensure a fair comparison between our results, mainly GO term enrichment results, Li et al. (2023) provided the list of genes tested for selection and those identified by BUSTED[S] as positively selected. We again used WebGestalt v. 2019 (Liao et al., 2019) to identify if genes with *d_N_*/*d_S_*≥1.00 were enriched gene ontology (GO) Biological Process and Cellular Component ontologies using the over-representation analysis (ORA) with the minimum number of genes for a category set to 5 (default) 30 (as in our analyses), the 618 genes with *d_N_*/*d_S_*≥1.00 as the foreground and the 4,444 genes tested for selection as the background gene set. These 618 genes were not enriched in any ontology after FDR correction; however, many terms were enriched with hypergeometric *P*≤0.05. None of these terms were the same as we identified (**Supplementary dataset 3**).

## Data availability

HyPhy json out files from ABSREL, BUSTED, and FEL (http://doi.org/10.5061/dryad.h44j0zpt4), alignments (http://doi.org/10.5061/dryad.5dv41nsbc), and the phylogenetic tree used for selection tests (http://doi.org/10.5061/dryad.j0zpc86n3) are available from Dryad. HyPhy json output files can be visualized with HyPhy Vision in any web browser: http://vision.hyphy.org and https://observablehq.com/@spond/busted-compare.

## Acknowledgments

This study was supported by NIH award 1R56AG071860-01 to V.J.L. Computational support was provided by the Center for Computational Research at the University at Buffalo. We thank S. L. Kosakovsky Pond for advice and help with running HyPhy, X. Zhou for clarifying the details of their analyses and providing IDs for their 4,444 gene dataset, and D. Enard for advice and thoughtful comments on this manuscript.

---

**Figure 1 – figure supplement 1.**
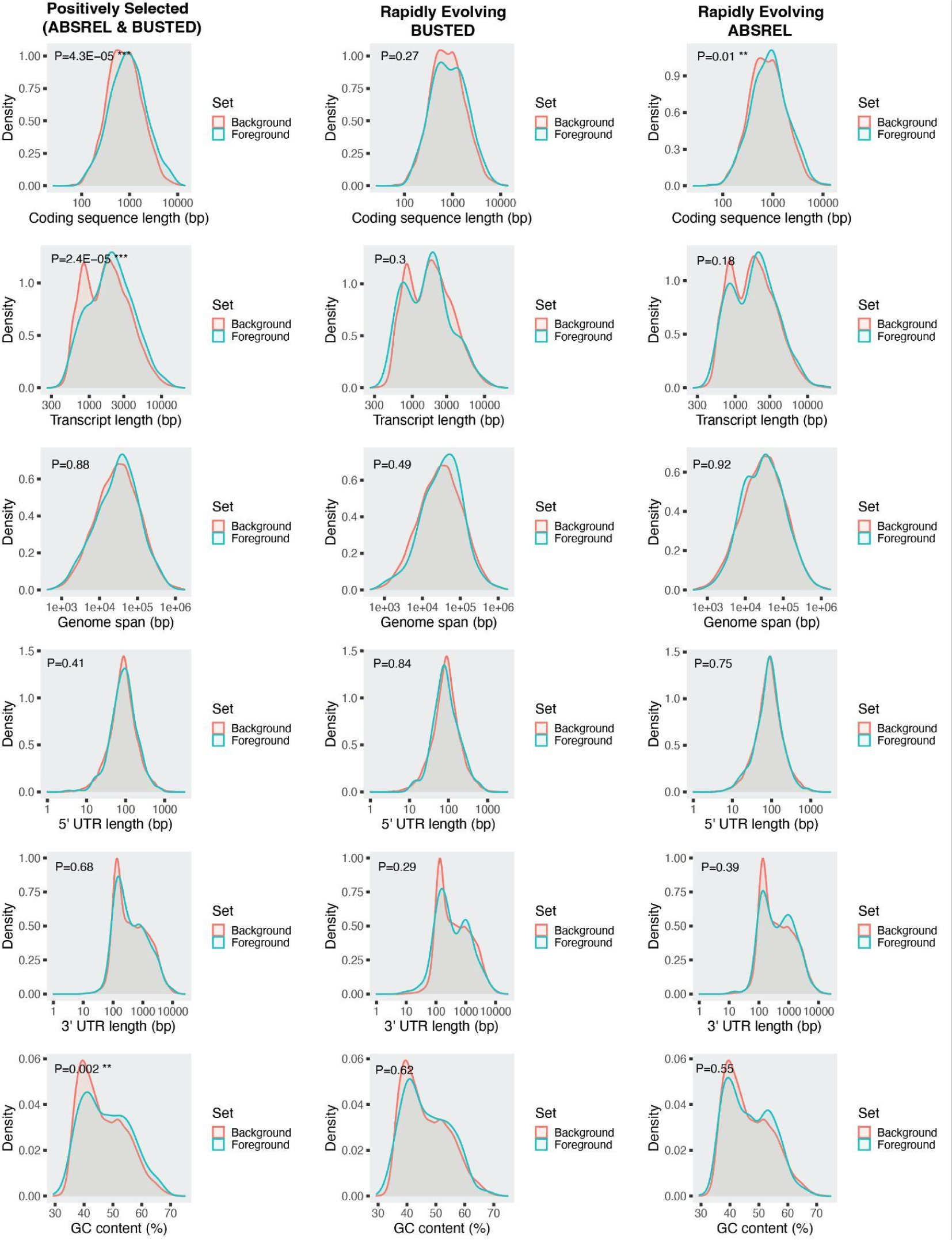
Genomic features of positively selected and rapidly evolving genes (foregroundf) compared to all genes tested (Background). *P*-values are derived from a Chi-square test.

