## Supplementary figures and images for "Rapid evolution of genes with anti-cancer functions during the origins of large bodies and cancer resistance in elephants"

### goslim_summary_wg_result1705335984.png

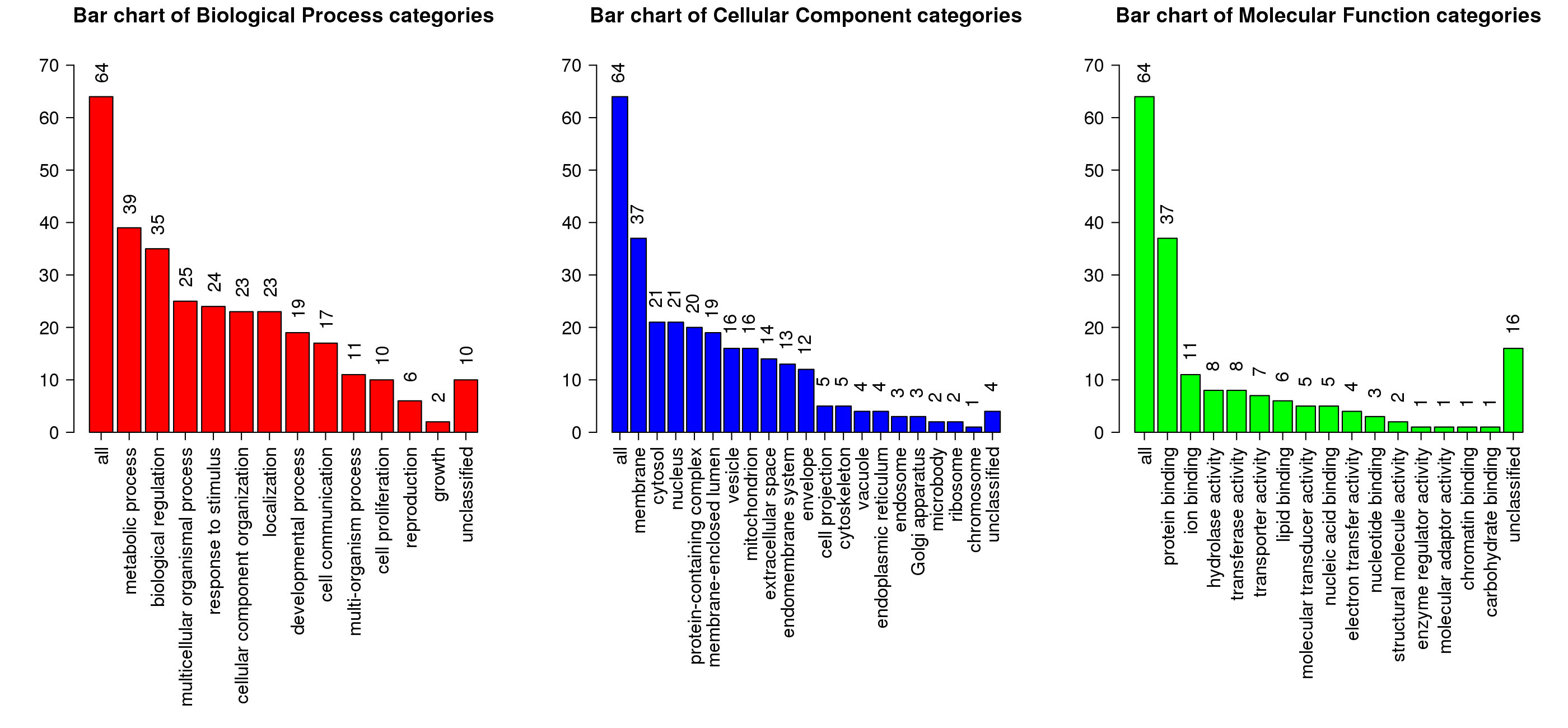

### goslim_summary_wg_result1705340001.png

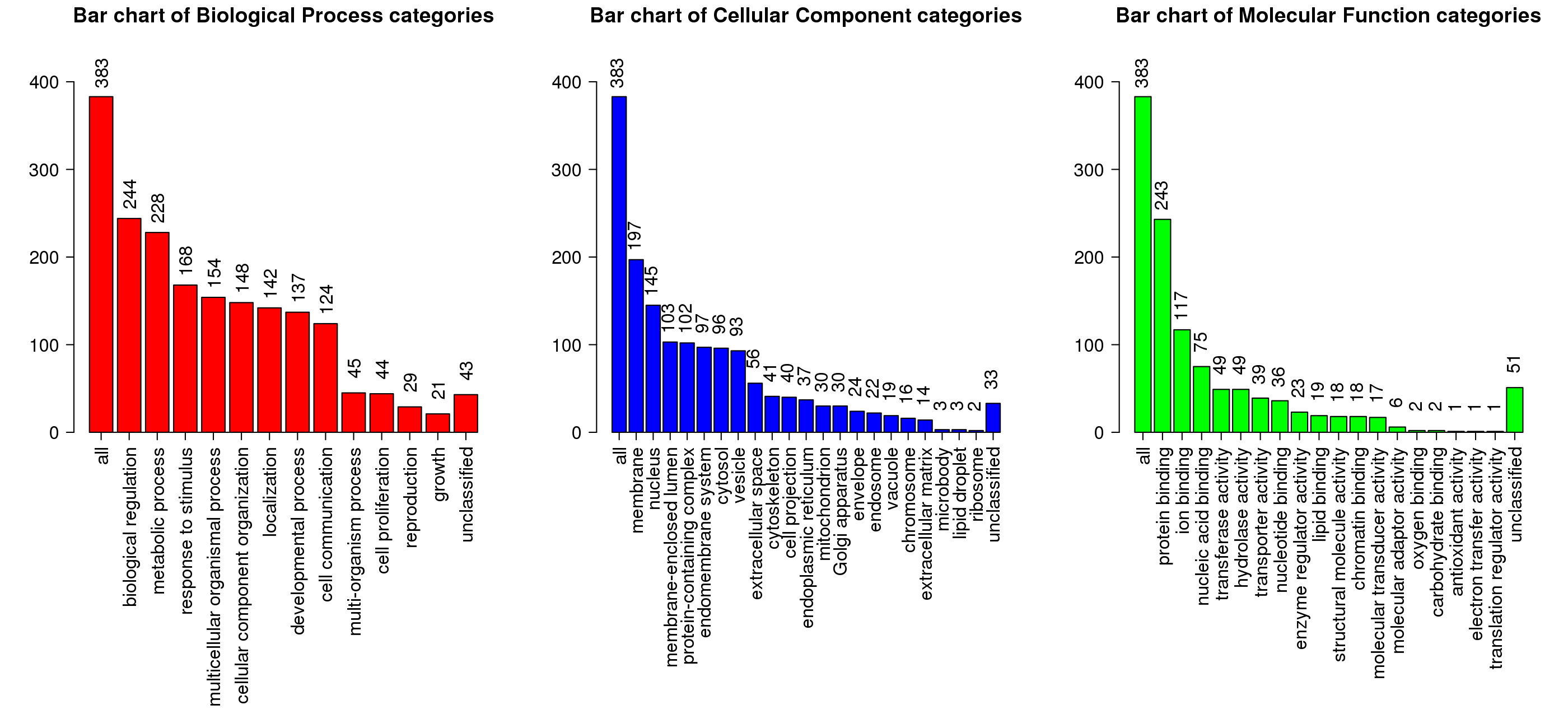

### goslim_summary_wg_result1707501092.png

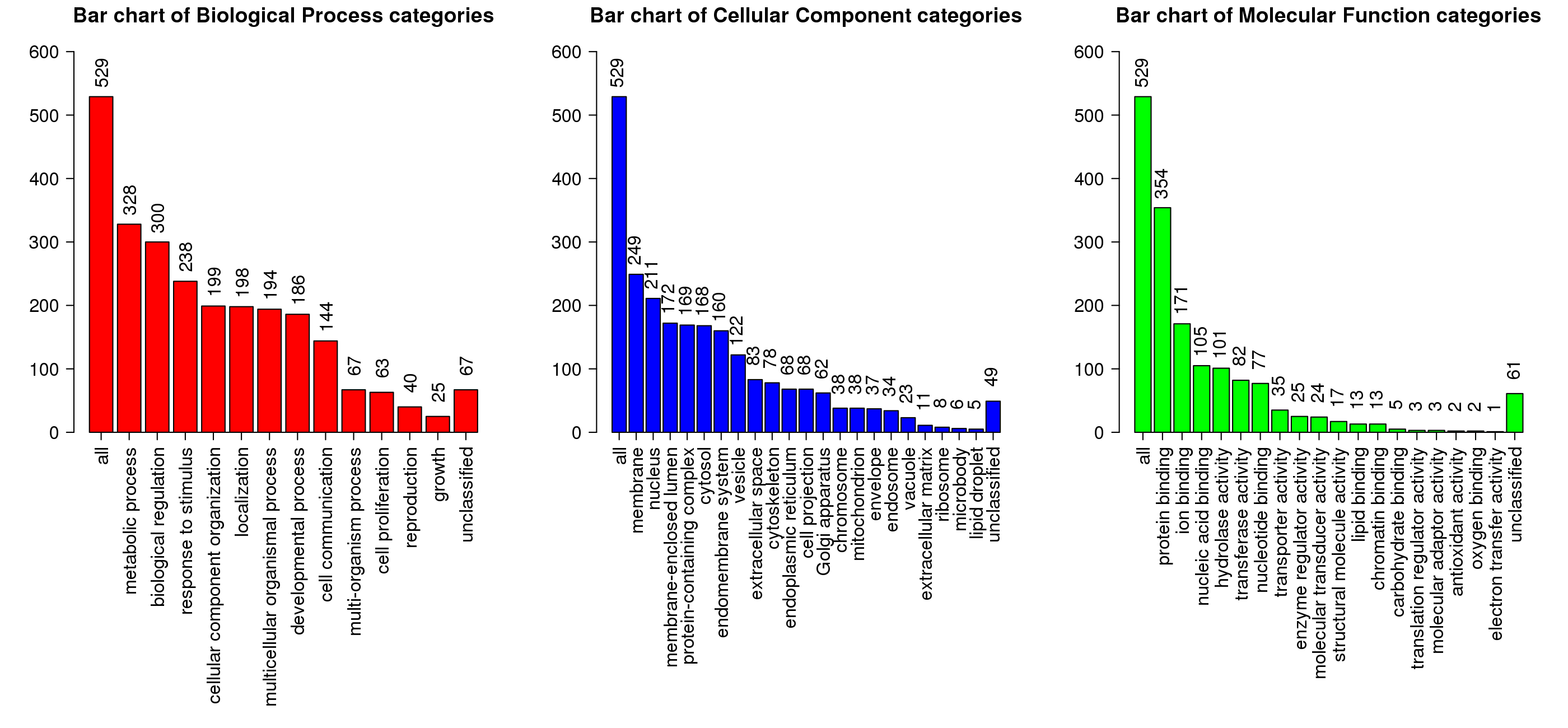

### goslim_summary_wg_result1708617015.png

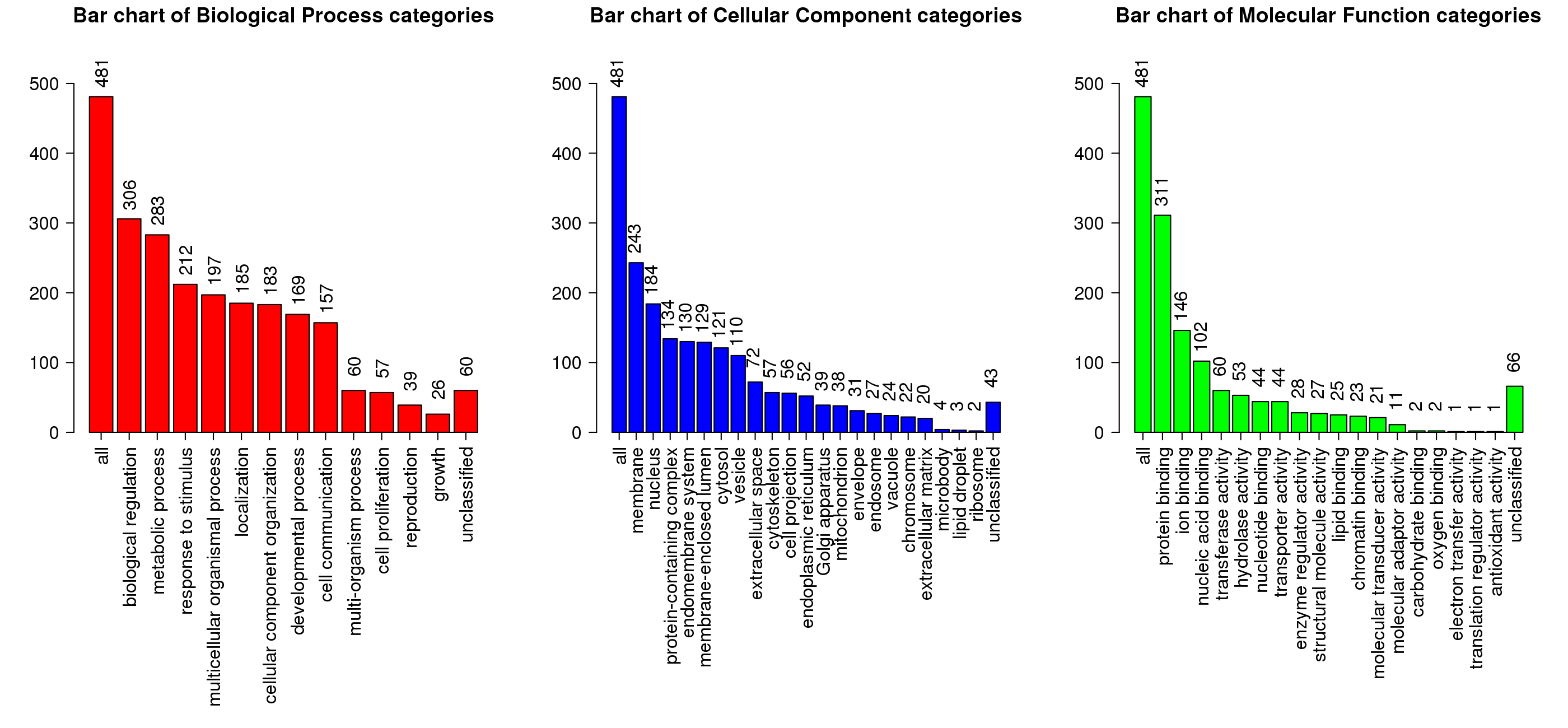

### goslim_summary_wg_result1708619288.png

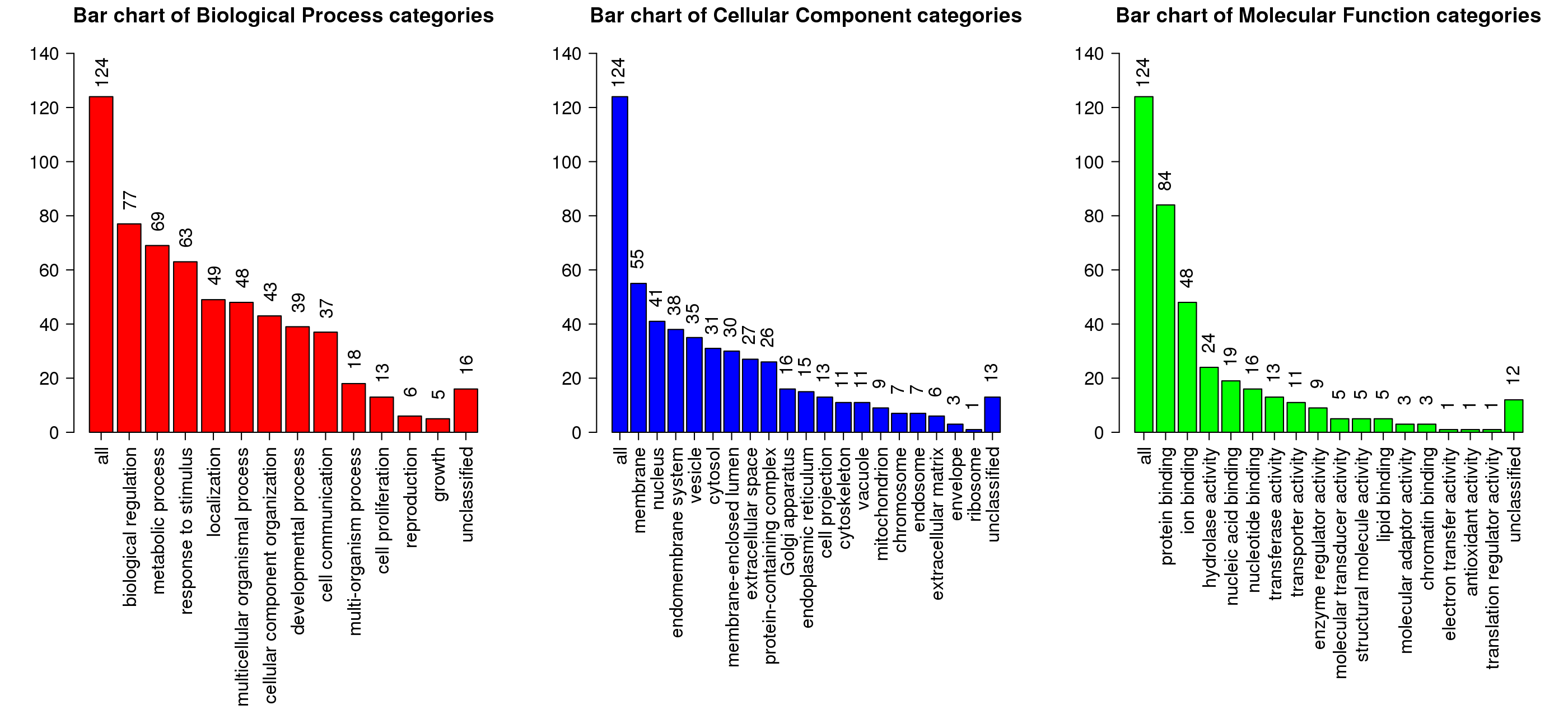

### goslim_summary_wg_result1708619336.png

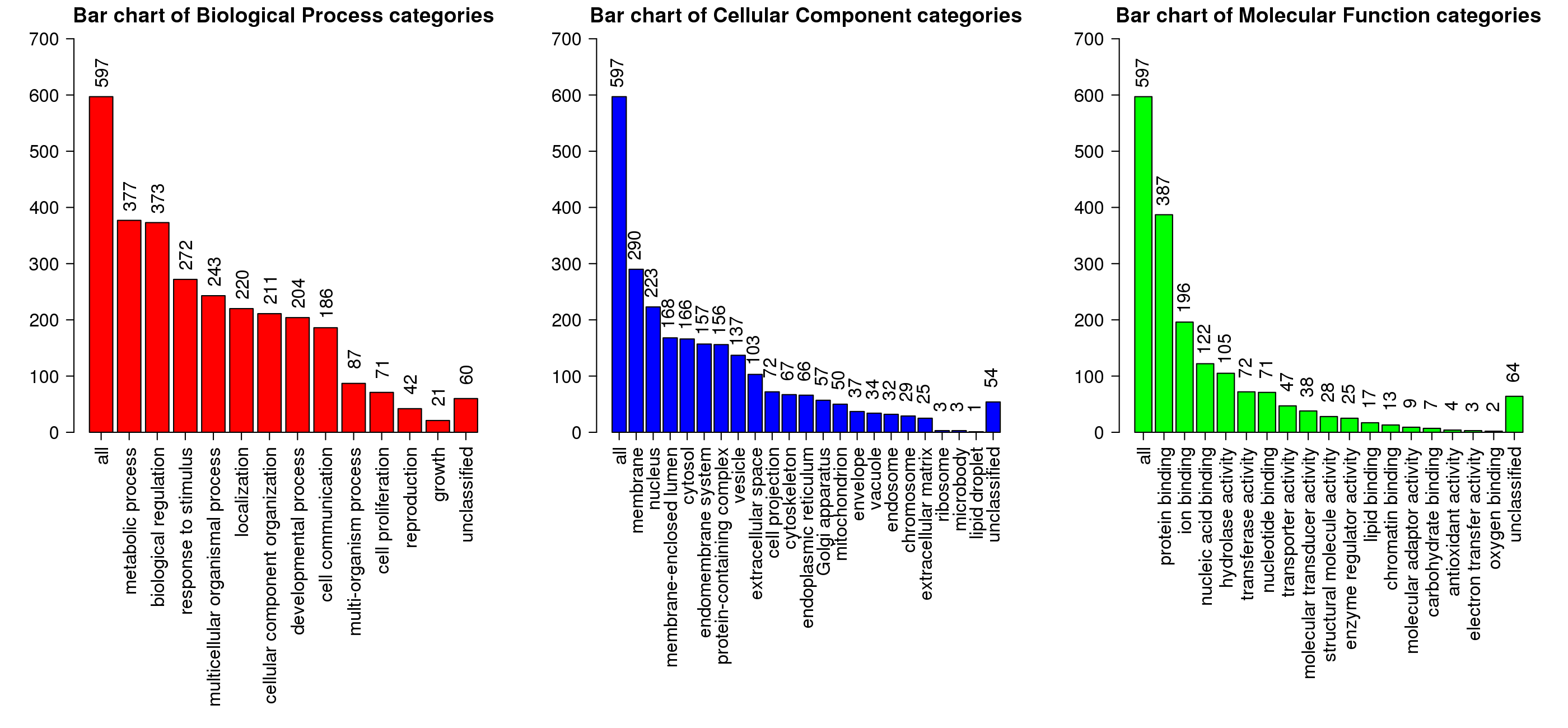
