## Supplementary material for "Rapid evolution of genes with anti-cancer functions during the origins of large bodies and cancer resistance in elephants": Figure 3 and 4 source data: Report_wg_result1708619288.html

WebGestalt (WEB-based GEne SeT AnaLysis Toolkit)


WEB-based GEne SeT AnaLysis Toolkit

Translating gene lists into biological insights...


---

#### Summary

Result Download

Job summary

- **Enrichment method:** ORA
- **Organism:** hsapiens
- **Enrichment Categories:** uploads/Cell Death Modes\_1708619288.gmt **ID Type:** genesymbol
- **Interesting list:** ABSREL\_best\_model\_says\_selection p=0.05-0.2\_1708619288.txt. **ID type:** genesymbol
- The interesting list contains **127** user IDs in which **124** user IDs are unambiguously mapped to **124** unique entrezgene IDs and **3** user IDs can not be mapped to any entrezgene ID.
- The GO Slim summary are based upon the **124** unique entrezgene IDs.
- Among **124** unique entrezgene IDs, **35** IDs are annotated to the selected functional categories and also in the reference list, which are used for the enrichment analysis.
- **Reference list:** uploads/GeneSet HUGO\_1708619288.txt **ID type:** genesymbol
- The reference list can be mapped to **13100** entrezgene IDs and  **13100** IDs are annotated to the selected functional categories that are used as the reference for the enrichment analysis.

**Parameters for the enrichment analysis:**

- **Minimum number of IDs in the category:** 5
- **Maximum number of IDs in the category:** 2000
- **FDR Method:** BH
- **Significance Level:** FDR < 0.1

Based on the above parameters, **3** categories are identified as enriched categories and all are shown in this report.

GO Slim summary for the user uploaded IDs

Each Biological Process, Cellular Component and Molecular Function category is represented by a red, blue and green bar, repectively.

The height of the bar represents the number of IDs in the user list and also in the category.

#### Enrichment Results

Redundancy reduction:
All


Affinity propagation


Weighted set cover


WebGestalt is currently developed and maintained by Yuxing Liao, Suhas Vasaikar, Zhiao Shi and Bing Zhang at the  Zhang Lab. Other people who have made significant contribution to the project include Jing Wang, Dexter Duncan, Stefan Kirov and Jay Snoddy.

**Funding credits:** NIH/NCI (U24 CA210954); Leidos (15X038); CPRIT (RR160027); NIH/NIAAA (U01 AA016662, U01 AA013512); NIH/NIDA (P01 DA015027); NIH/NIMH (P50 MH078028, P50 MH096972); NIH/NCI (U24 CA159988); NIH/NIGMS (R01 GM088822).
